# Experimental short-term heatwaves negatively impact body weight gain and survival during larval development in a wild pollinator

**DOI:** 10.1101/2024.10.09.617416

**Authors:** Laura Wögler, Christoph Kurze

## Abstract

Climate change-induced heatwaves threaten global biodiversity, including crucial pollinators like bumblebees. Despite alarming projections, little is known about the effects of short-term heatwaves on insect larval development. Hence, we investigated the impact of simulated heatwaves on the development of fourth instar larvae (L4) of *Bombus terrestris* L. (Hymenoptera: Apidae) using an *in vitro* rearing method. Individual larvae were incubated at 37°C and 38°C for a period of 4 days, with a constant rearing temperature of 34°C as the control. We examined body weight gain, developmental duration, survival to adult stage, and adult body size (i.e. dry mass, ITD, and head width). A simulated heatwave of 37°C did not significantly affect larval development, but 38°C impaired larval body mass gain. While developmental duration and adult body size were unaffected, an acute heat stress of 38°C during the L4 stage reduced the probability of pupae reaching adulthood. These findings highlight the potential for heatwaves to negatively affect bee populations by impairing larval growth and reducing survival to the adult stage, which may have severe implication for colony fitness.

## 1. Introduction

Climate change is a major challenge of the 21^st^ century, causing cascading effects that impact weather patterns, biodiversity, and entire ecosystems (Garcia *et al*. 2014; IPCC 2023). Particularly concerning are heatwaves, defined as at least three consecutive days with temperatures exceeding a certain threshold, which have increased in frequency and intensity (Lhotka, Kyselý & Farda 2018; Stillman 2019; Perkins-Kirkpatrick & Lewis 2020). These heatwaves have a significant impact on terrestrial animals, including humans (Zuo *et al*. 2015; Stillman 2019), small mammals (Ratnayake *et al*. 2019; Zhao *et al*. 2020; Fuller *et al*. 2021), birds (McKechnie & Wolf 2010; Conradie *et al*. 2019), and insects (González-Tokman *et al*. 2020; Ma, Ma & Pincebourde 2021; Bodlah *et al*. 2023). Despite this threat, our understanding of the impact of heatwaves on animal development, their fitness, and populations remains limited (Jentsch, Kreyling & Beierkuhnlein 2007; Stillman 2019; González-Tokman *et al*. 2020; Fuller *et al*. 2021; Ma, Ma & Pincebourde 2021).

Cold-adapted heterothermic species like bumblebees (*Bombus* sp.), which have species-specific distributions ranging from the Arctic to temperate/Mediterranean regions, may be particularly at risk (Rasmont & Iserbyt 2012; Soroye, Newbold & Kerr 2020; Suzuki-Ohno *et al*. 2020; Maebe *et al*. 2021; Martinet *et al*. 2021a; Ghisbain *et al*. 2024). As key pollinators in many ecosystems and for agriculture (Corbet, Williams & Osborne 1991; Klein *et al*. 2007; Potts *et al*. 2016; Cameron & Sadd 2020), understanding how extreme heat events impact their physiology and fitness is crucial. Recent studies show that heat stress can reduce adult survival (Kuo *et al*. 2023; Quinlan *et al*. 2023) and elevated temperatures may lower colony fitness (Martinet *et al*. 2021b; Theodorou *et al*. 2022). Under heatwave-like temperatures, adult bumblebees exhibit impaired cognition (Gérard *et al*. 2022a) and scent perception (Nooten *et al*. 2024), and display altered fanning and foraging behaviours (Kuo *et al*. 2023; Bretzlaff, Kerr & Darveau 2024; Sepúlveda *et al*. 2024), potentially affecting colony fitness. Interestingly, foraging behaviour and response to stimuli are altered in adults even when exposed to heatwave-like temperatures during their larval and pupal development (Gérard *et al*. 2022b; Perl *et al*. 2022). Exposing entire colonies to high temperatures for extended periods can lead to alterations in wing size asymmetry, wing shape and size, and reductions in body and antennae sizes (Gérard *et al*. 2018; Guiraud *et al*. 2021; Gérard *et al*. 2022b; Perl *et al*. 2022; Gérard *et al*. 2023). Despite that different developmental stages are likely to be variably affected, it is unclear which stages are particularly vulnerable to heatwaves and whether shorter extreme heat events are sufficient to impair their development.

To address this knowledge gap, we adapted an *in vitro* rearing protocol (Kato *et al*. 2022) to examine the direct impact of a four-day long heat stress during the development in 4^th^ instar larvae (L4). This experimental approach prevented the mitigation of heat through worker fanning activities (Weidenmüller, Kleineidam & Tautz 2002). While *in vitro* rearing is a standard procedure in honeybee research (Crailsheim *et al*. 2013; Schmehl *et al*. 2016), it is rarely used in bumblebee research (Pereboom, Velthuis & Duchateau 2003; Kato *et al*. 2022). Therefore, we chose *B. terrestris* as a model species, although it is known to be rather heat-tolerant (Zambra *et al*. 2020; Martinet *et al*. 2021a). We used this method, to investigate changes in body mass, developmental duration, survival until pupation and emergence, and adult body size following a four-day long heatwave-like exposure during L4 development.

## 2. Methods

### (a) Experimental overview

To simulate the effect of short-term heatwaves on larval development, we collected L4 larvae (total of 289) from five commercial *Bombus terrestris* colonies to rear them *in vitro*. These larvae were pseudo-randomly assigned to one of the three experimental groups to ensure equal distribution among treatments and colonies. Larvae of the simulated heatwave treatment were exposed to either 37°C (n = 96) or 38°C (n = 92) for four days, while the control group (ctrl, n = 101) was reared at a constant 34°C (Pereboom, Velthuis & Duchateau 2003; Kato *et al*. 2022). We recorded body mass changes during treatment and papal stage. Their survival was checked daily until adult bees emerged and reached the age of 2 days. These adult bees were freeze killed and kept at -20°C for subsequent morphometric measurements and analysis of their dry mass and lipid content.

### (b) Colony husbandry

Upon arrival each *Bombus terrestris* colony (Natupol Research Hives, Koppert B.V., Netherlands) consisted of 20-30 workers, brood, and one queen. They were housed and maintained under similar conditions as described previously (Gilgenreiner & Kurze 2024). Briefly, bumblebees had access to 70% (w/v) sucrose solution *ad libitum* in a foraging arena (59 (l) x 39 (w) x 26 (h) cm) with 14:10 h light:dark regime. Depending on colony size, each colony received 6-11 g of pollen candy daily. The room temperature was maintained at 25 °C ± 1 °C and 30-50% relative humidity. We allowed all colonies to develop for at least 1 month before starting to collect L4 larvae for the experiment.

### (c) Collection of 4^th^ instar larvae

Before collecting L4 larvae for our experiment, we carefully removed all existing L4 larvae from the colonies using soft tweezers. This allowed us to identify and collect larvae that just entered the L4 stage during our daily colonies checks over the following days. L4 larvae are easily identified by their individual, spherical brood cells with a small opening for food provisioning (Tian & Hines 2018). To facilitate the collection process, we temporarily moved all adult bumblebee to a separate cage and returned them afterwards.

### (d) *In vitro* rearing

We followed established *in vitro* rearing procedures with slight modifications (Crailsheim *et al*. 2013; Kato *et al*. 2022), where we carefully transferred L4 larvae into 3D-printed polylactide (PLA) artificial brood cells (capacity: 0.6 ml, Ø 8 mm, figure 1a). This facilitated measuring their weight gain without touching them again. To simulate short-term heatwaves, we randomly assigned L4 larvae to one of the three experimental groups and either reared them at 37°C and 38°C for 4 days (KB115, BINDER GmbH, Germany). The control group (ctrl) was maintained at constant temperature of 34°C (Pereboom, Velthuis & Duchateau 2003; Kato *et al*. 2022). Those artificial brood cells were placed into 24-or 48-well clear flat bottom plates (Falcon/Corning, USA) (figure 1b) and kept inside in ventilated plastic containers (18.5 × 18.5 × 11.5 cm) together with a 120 mL cup of saturated sodium chloride solution to maintain 65 ± 10 % relative humidity.

**Figure 1.**
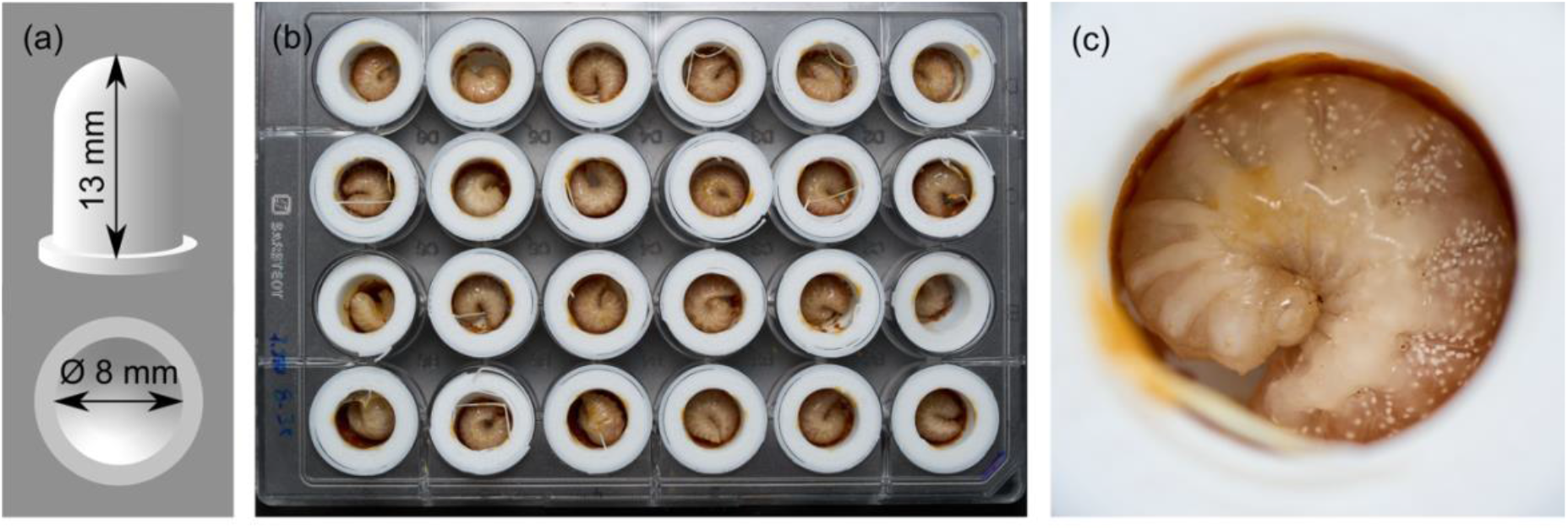
*In vitro* rearing of L4 larvae. (a) Design of 3D-printed polylactide (PLA) artificial brood cells (capacity: 0.6 ml), (b) artificial brood cells with L4 larvae inside a 24-well clear flat bottom plate, and (c) L4 feeding on medium.

Larvae were fed with a pollen medium twice daily, in the morning and in the evening. The medium contained 50% w/v sucrose solution (Südzucker AG, Germany), 40% honeybee collected organic pollen (Bio-Blütenpollen, naturwaren-niederrhein GmbH, Germany), 10 % Bacto yeast extract (Bacto™, BD, USA), and 1 % casein sodium salt from bovine milk (Sigma-Aldrich, Germany). Aliquots of medium were stored at -20°C and warmed up to 34°C and vortexed before feeding. A feeding session consisted of two 20-minute rounds on a heated plate at 35°C (Medax model 12801, Medax Nagel GmbH, Germany). Larvae were initially fed a 6 μL droplet (7.1 ± 1.6 mg) of medium onto their ventral abdomen (figure 1c). We monitored larval behaviour to determine satiation. A larva was considered satiated when it curled up and ceased movement (Supplementary video 1) and hungry when it remained active. Larvae that did not consume food during the first feeding round were not offered additional food. At the end of the feeding session, any remaining food was carefully removed to prevent the larvae from suffocating due to dried food blocking their trachea. Larvae entering pupal stage were no longer fed.

### (e) Measurements of body mass

Each individual bee was weight at four different stages: as L4 larvae pre- and post-treatment, at the beginning of pupal stage, and as a newly emerged adult using a fine scale (d = 0.1 mg, Sartorius AC120S, Sartorius AG, Germany). To reduce stress and avoid any potential handling damage of the larvae, we kept them inside their 3D-printed cell. Consequently, we subtracted the empty cell weight to obtain the actual weights.

### (f) Morphometric measurements, dry mass, and lipid content

The intertegular distance (ITD) and dry mass serve as a proxy for adult workers’ body size (Kendall *et al*. 2019; Gilgenreiner & Kurze 2024). In addition to the ITD, we also measured the head width using a digital microscope (CHX-500F, Keyence GmbH, Germany). We obtained individual dry mass and lipid content following our previous protocol (Gilgenreiner & Kurze 2024). Briefly, we dissected the ventral abdominal segments and dried corpses at 60°C for 3 days in a drying cabinet (U40, Memmert GmbH & Co. KG, Germany). After weighing their dry mass (d = 0.1 mg, analytic balance M-Pact AX224, Sartorius GmbH, Germany), we extracted their body lipid with petroleum ether for 5 days. After discarding the ether and rinsing them with fresh ether, the bees were dried for an additional 3 days and weighed again. The lipid contented was calculated as the difference between the initial and post-extraction dry weights.

### (g) Statistical analyses

All statistical analyses and data visualizations were performed using R version 4.4.1 (R-Team 2018). The complete code and output are provided in the electronic supplementary material. Briefly, the probability of L4 larvae reaching papal stage and adulthood was calculated using *chisq-test* function to perform a Pearson’s *χ*^2^ test for count data with Yate’s continuity correction. Pairwise comparisons of survival probabilities between survival treatment groups were conducted using the *pairwise*.*prop*.*test* function with Benjamini-Hochberg correction. In addition, we ran generalized linear mixed effect models (GLMMs) using the *glmmTMB* package with an gaussian data distribution to analyse effects of the heatwave treatment as a fixed factor on the duration to pupation and to emergence, the relative body weight gain during treatment, the relative body weight loss during pupation, adult body size (with dry mass, ITD, and head width serving as proxies), and the relative lipid content as response variables. In addition, we ran another GLMM based on binomial data distribution to test whether both heatwave and relative body weight gain as well as their interactions affected their probability to reach pupal stage and adulthood. We included colony ID as a random factor in all models to account for colony-specific covariates. Model selection was performed based on the Akaike information criterion (AIC) and likelihood ratio tests. The final models were compared with their respective null-models. Model assumptions and dispersion of the data were checked using the *DHARMa* package (Hartig 2020). As model assumptions were not met when analysing treatment effects on relative body weight loss during pupation, we used a Kruskal-Wallis rank sum test. Significance levels (p < 0.05) were determined using the *Anova* function of the *car* package (Zuur *et al*. 2009). Pairwise comparisons between treatment groups were conducted using the function *emmeans* (Lenth & Lenth 2018) adjusted with the Tukey’s HSD.

## 3. Results

While there was no significant difference in the number of L4 larvae reaching the pupal stage (*χ*^2^ = 1.36, df = 2, p = 0.507; figure 2a), a four-day-long heatwave at the beginning of L4 development significantly impacted their probability of reaching adulthood (*χ*^2^ = 6.79, df = 2, p = 0.034; figure 2c). Larvae exposed to 38°C were 50% less probable to reach adulthood compared to larvae exposed to 37°C (p = 0.024) or the control group (p = 0.024). However, there was no significant difference between the 37°C and the control group (p = 1). There was also no significant effect of heatwave treatment on the developmental times for larvae reaching pupal stage (glmm: *χ*^2^ = 3.28, df = 2, p = 0.194; figure 2b) and for pupae until emergence (*χ*^2^ = 0.66, df = 2, p = 0.718; figure 2d).

**Figure 2.**
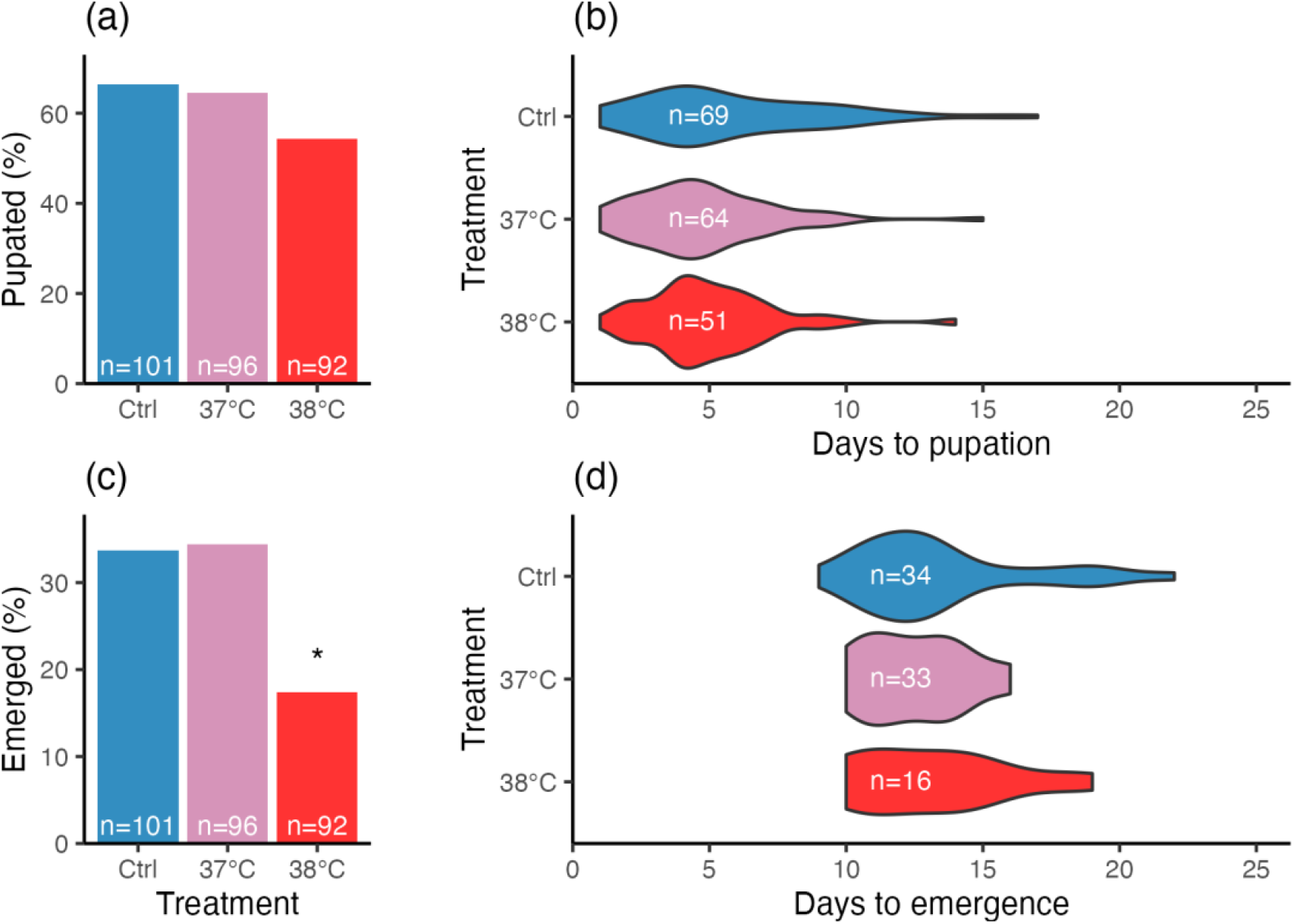
Effects of 4-day-long simulated heatwaves on survival and developmental duration of L4 larvae (a, c) and pupae (c, d). The controls (Ctrl, reared at constant 34°C, in blue) were compared to individuals that had been exposed to increased rearing temperatures at 37°C (in rose) and 38 °C (in red) during L4 development for 4 days. Significant differences (p < 0.05) are denoted with an asterisk (*).

Heatwaves significantly affected the relative weight gain in L4 larvae during the treatment (glmm: *χ*^2^ = 7.94, df = 2, p = 0.019; figure 3a). While an exposure to 38°C revealed significant lower weight gains compared to the control (Tukey HSD: t-ratio = 2.80, p = 0.015), there was no significant effect for larvae exposure to 37°C (t-ratio = 1.61, p = 0.242). Regardless of treatment, their body weight loss during pupation had no significantly affected (Kraskal-Wallis *χ*^2^ = 3.80, df = 2, p = 0.150; figure 3b). There were also no significant effects on morphometrics in 2-day-old adults, including their dry mass (glmm: *χ*^2^ = 2.52, df = 2, p = 0.284; figure 3c), ITD (*χ*^2^ = 2.834, df = 2, p = 0.243; figure 3d), heat width (*χ*^2^ =, df =, p = 0.; figure 3e), and relative lipid content (*χ*^2^ =, df =, p = 0.; figure 3f). However, it is worth noting that the sample size was significantly reduced in the 38°C treatment group at the adult stage (n = 16, see figure 2c,d).

**Figure 3.**
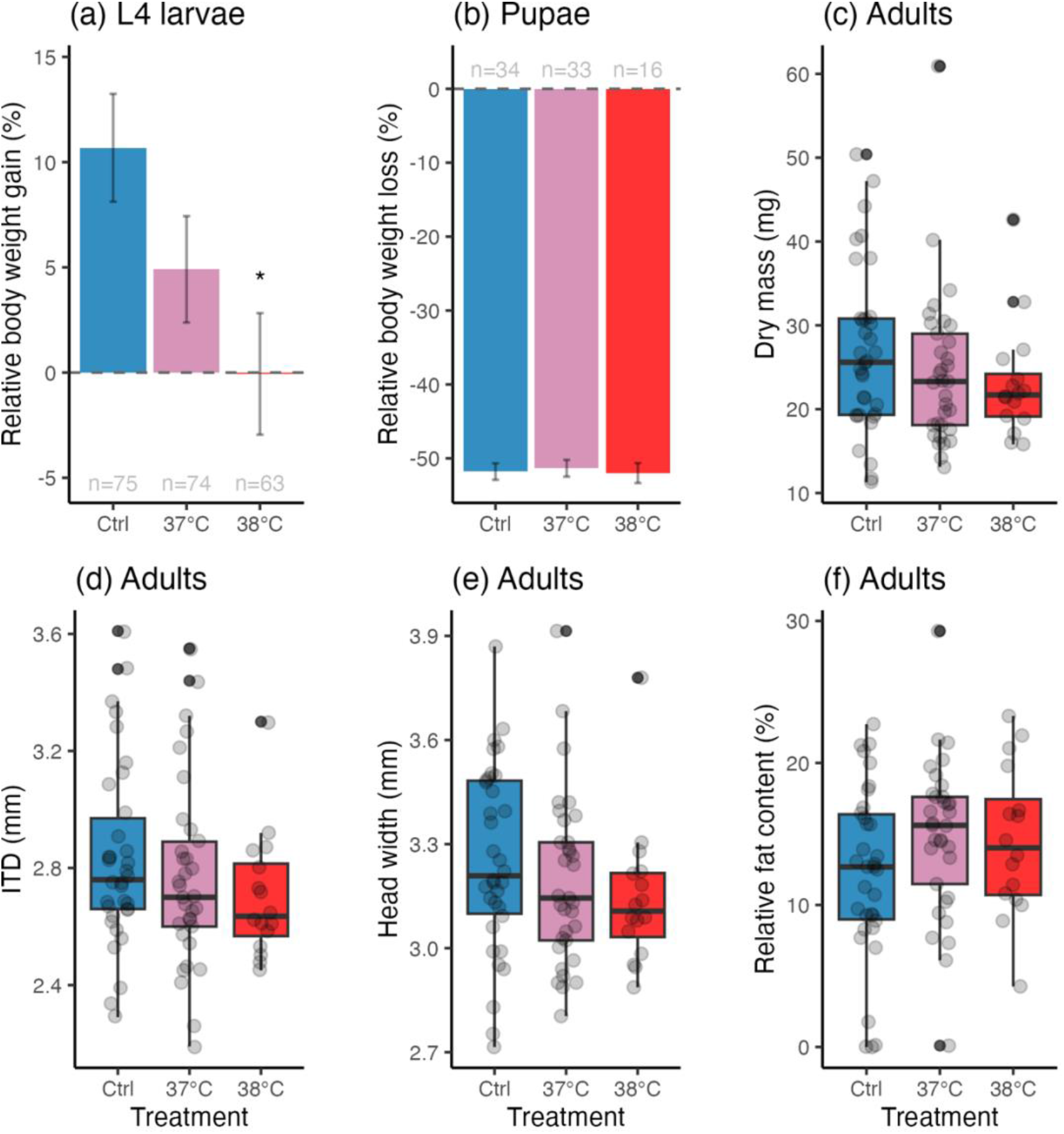
Effects of 4-day-long simulated heatwaves during L4 stage on bumblebee morphometrics: (a) relative body mass gain during treatment (means ± s.e.); (b) relative body mass loss during pupation (means ± s.e.); and (c-f) consequences on their dry mass, ITD (intertegular distance), head width, and relative lipid content as 2-day-old adults. The controls (Ctrl, reared at constant 34°C, in blue) were compared to individuals that had been exposed to increased rearing temperatures at 37°C (in rose) and 38 °C (in red) during L4 development for 4 days. Significant differences (p < 0.05) are denoted with an asterisk (*).

We found that both the heatwave treatment and relative body weight gain during treatment impacted the probability of larvae to reach adulthood (treatment: *χ*^2^ = 14.759, df = 2, p = 0.001; weight gain: *χ*^2^ = 24.35, df = 1, p < 0.0001), but not their interactions (*χ*^2^ = 0.04, df = 2, p = 0.980). In addition, we found that relative body weight gain had a significant effect on the probability of larvae to pupate (*χ*^2^ = 14.10, df = 1, p = 0.0002).

## 4. Discussion

Our data provides evidence of how simulated short-term heatwaves during the L4 stage affects development and survival until reaching adulthood in *B. terrestris* (figures 1c and 2a). Nonetheless, there was no effect on the duration of larval development (figure 2b,d) nor on the morphometrics in adults (figure 3c-e). This is in contrast to previous studies showing that exposing colonies to elevated temperatures would produce smaller workers, indicated by smaller ITD (Guiraud *et al*. 2021; Gérard *et al*. 2023), with reduced antennae in *B. terrestris* (Gérard *et al*. 2023). In addition, while wing size and shape can be affected (Gérard *et al*. 2018), this may not always be the case (Gérard *et al*. 2023). As only 17% of individuals emerged as adults in the 38°C treatment group of our experiment, this sample size was too small to thoroughly analyse wing morphology. Regardless, a potential explanation for this discrepancy could be that colonies were exposed to higher temperatures for extended periods in those experiments, whereas we tested the effect of short-term heatwave-like exposures in L4 larvae. This explanation would be supported by another study showing that body or organ sizes were also not altered when exposing colonies to elevated temperatures for shorter periods (Perl *et al*. 2022).

We found a 50% lower probability to emerge as adults of L4 larvae exposed to 38°C compared to both the 37°C heatwave and the control group (figure 2c). Although the emergence rate of our *in vitro* reaching was low, our pupation rate, ranging between 55-66% irrespective of treatment, was similar to previous research (Kato *et al*. 2022). This suggests that 38°C might be a threshold temperature with ripple effects on critical processes during pupation. By contrast, constantly elevated temperatures were experimentally shown to increase drone production in microcolonies of *B. terrestris* up to nest temperatures of 34-36°C, at which workers dramatically spend more time with fanning (Sepúlveda *et al*. 2024). In *B. impatiens*, offspring production already decrease at 35°C as workers abandon their colony (Bretzlaff, Kerr & Darveau 2024). In our experiment, we took larvae out of their natural nest environment and reared them *in vitro* without workers that could feed and incubate or cool L4 larvae. This gave us more experimental control over timing and control of the heatwave exposure. While our temperatures seem quite high, we think these are not unrealistic heatwave scenarios (Lhotka, Kyselý & Farda 2018; Stillman 2019; Perkins-Kirkpatrick & Lewis 2020) and are still comparable to previous studies, considering that the actual brood temperature is typically 2°C warmer than the ambient nest temperature (Vogt 1986; Weidenmüller, Kleineidam & Tautz 2002).

Our *in vitro* rearing approach also allowed us to closely monitor weight gain and loss throughout larvae development until emergence (figure 3a-b). These data revealed that a 4-day-long exposure to 38°C resulted in lower weight gains during the L4 development compared to both the 37°C heatwave and control group (figure 3a), indicating that those larvae consumed less food, potentially due to reduced appetite. While it is known that stress reduces food intake and consequently weight gain in mammals (Rabasa & Dickson 2016), little is known about how acute stress affects food consumption and body weight gain in insects. Reaching a critical weight during the larval development, however, is known to be crucial to initiate molting and metamorphosis as shown in the tobacco hornworm (*Manduca sexta* L., Lepidoptera: Sphingidae) (Nijhout & Williams 1974). Our data confirms this for bumblebees, as the weight gain at L4 stage significant predictor for pupation success irrespective of the treatment group. In fruit flies (*Drosophila melanogaster* M., Diptera: Drosophilidae) short-term heat stress of 24 h in 3^rd^ instar larvae reduced food intake in adults on their first day after emergence without impacting their body weight but leading to increased glucose and trehalose levels while reducing lipid stores (Karpova *et al*. 2024). Likewise, we did not find any difference in dry body mass in 2-day-old adult bees across treatment groups (figure 3c), but also not in the relative lipid content (figure 3f). One potential explanation could be that our extreme heat stress may have allowed that individuals with adaptive advantages to emerge as an adult, without affecting their overall development.

Although both the heatwave treatment and the weight gain had a significant effect on the probability of adults reaching adulthood, there was no interaction between both factors, suggesting that weight gain did not increase the survival outcome in the 38°C heatwave group. Regardless, our data supports the hypothesis that acute stress during larval development has a drastic impact at a later life stage (Karpova *et al*. 2024), although effects could have also just been delayed in our experiment. An exposure to 38°C at the L4 stage marked a threshold at which processes during metamorphosis are likely impaired in *B. terrestris*. It would be interesting to see how acute stress in early development would impact the life-history of adult bees. It has already been shown that heatwave-like temperatures during late development impact initial behavioural responses to sensory stimuli in adult workers of *B. terrestris* (Perl *et al*. 2022). This could not only have detrimental effects for the individual worker but also ripple effects on colony fitness.

In conclusion, while our *in-vitro* rearing experiment showed a certain resilience of *B. terrestris* larvae to heatwave-like exposures up to 37°C, extreme temperatures of 38°C appeared to be the threshold where pupal development was severely impaired. Individuals reaching adulthood, however, did not differ in their body size (dry mass, ITD, and head width) and relative lipid content, suggesting potential adaptive advantages in those surviving bees. With our experimental approach we aimed to investigate the specific effects of acute thermal stress at the L4 stage, which we traded for experimental realism. Larvae were taken out of their natural nest environment which likely has a large impact on their survival. Although our selected temperatures may appear extreme, they are only 3°C and 4°C above what is considered to optimal rearing conditions. Thus, we think that our scenarios are not totally unrealistic. Furthermore, we studied *B. terrestris* as a model species, which is rather heat-tolerant. We would expect more severe effects in more cold-adapted species such as *B. lapidarius, B. alpinus* or *B. poralris*. Given the increasing frequency and severity of heatwaves, it is crucial to investigate their impact on the life-history and adaptive potential of keystone species like bumblebees.

## Data accessibility

All data and statistical code output supporting the findings of this study are provided in the electronic supplementary material.

## Acknowledgments

We thank Franziska Altmann, Erik Köhler, Sandra Laußer and Maximilian M. Mandlinger for their assistance in the laboratory. We would like to extend our gratitude to Dr. Tomer J. Czaczkes for sposoring the bumblebee colonies.

## Author’s contributions

LW carried out the experimental work and gave approval for publication. CK conceived, designed, and supervised the study; analysed and interpreted the data; and wrote the manuscript.

## Ethics statement

This study was conducted in accordance with the ethical regulations of the German Animal Welfare Act (TierSchG) for conducting experiments with insects.

## Funding

This study was carried out without any third-party funding.

## Competing interests

The authors have no competing interests.

## Notes

### Competing Interest Statement

The authors have declared no competing interest.

